# Protein biocargo of citrus fruit-derived vesicles reveals heterogeneous transport and extracellular vesicles populations

**DOI:** 10.1101/321240

**Authors:** Gabriella Pocsfalvi, Lilla Turiák, Alfredo Ambrosone, Pasquale del Gaudio, Gina Puska, Immacolata Fiume, Teresa Silvestre, Károly Vékey

**Affiliations:** Institute of Biosciences and BioResources, National Research Council of Italy, via P, Castellino 111 - 80131 Naples, Italy; MS Proteomics Research Group, Institute of Organic Chemistry, Research Centre for Natural Sciences, Hungarian Academy of Sciences, Magyar tudósok körútja 2. Budapest, H-1117 Hungary; Department of Pharmacy, University of Salerno, Via Giovanni Paolo II, 132 - 84084 Fisciano (SA), Italy; Department of Anatomy, Cell and Developmental Biology, Eötvös Loránd University, Pazmany s. 1/C. 6.520. Budapest, H-1117 Hungary

**Keywords:** nanovectors, citrus, fruit juice sac cell, vesicles, extracellular vesicles, vesicular trafficking, proteomics, enzymes, oxidoreductases, hydrolases, bioinformatics

## Abstract

Cellular vesicles are membrane-enclosed organelles that transport material inside and outside the cell. Plant-derived vesicles are receiving increasing attention due to their potential as nanovectors for the delivery of biologically active substances. We aimed to expand our understanding about the heterogeneity and the protein biocargo of citrus fruit juice sac cell-derived vesicles. Micro- and nanosized vesicle fractions were isolated from four citrus species, *C. sinensis*, *C. limon*, *C. paradisi* and *C. aurantium*, characterized using physicochemical methods and protein cargos were compared using label-free quantitative shotgun proteomics. In each sample approximately 600-800 proteins were identified. Orthologues of most of the top-ranking proteins have previously been reported in extracellular vesicles of mammalian origin. Patellin-3-like, clathrin heavy chain, heat shock proteins, 14-3-3 protein, glyceraldehyde-3-phosphate dehydrogenase and fructose-bisphosphate aldolase 6 were highly expressed in all citrus vesicle fractions. The presence of membrane channel aquaporin on the other hand characteristic of the nanovesicle fractions. Bioinformatics revealed more than hundred protein orthologues potentially implicated in vesicular trafficking. In particular, CCV, COPI and COPII coat proteins indicates the presence of highly heterogeneous populations of intracellular transport vesicles. Moreover, the different hydrolases and oxidoreductases transported within the citrus fruit-derived vesicles can be responsible for the various biological activities possessed by the preparations.

**Abbreviations:**

EVs: extracellular vesicles;
MVs: microvesicles;
NVs: nanovesicles;
PM: plasma membrane;
UC: ultracentrifugation;
CCV: clathrin coated vesicles;
COPI: coat protein I;
COPII: coat protein II

## 1. Introduction

Cellular vesicles are fluid filled lipid bilayer enclosed organelles within or outside the plasma membrane (PM) used to transport materials. Formation and secretion of vesicles are evolutionary conserved and hence vesicles are ubiquitous in all three primary kingdoms, bacteria, archaea and eukaryotes. Vesicles can be classified based on their size, function, location and the nature of their cargo. In plant cells principally four types of vesicles can be distinguished: the large central vacuole (Echeverría, 2000), transport vesicles (Hwang and Robinson, 2009), secretory vesicles (van de Meene et al., 2017) and apoplastic vesicles (Rutter and Innes, 2017). The vacuole, transport and secretory vesicles are found intracellularly while apoplastic vesicles are extracellular. Plant vacuole plays an important role in molecular storage, degradation and maintenance of turgor pressure. Transport vesicles are involved in the exchange of lipids, nucleic acids, proteins and small soluble molecules between cellular organelles and compartments within the cytoplasm. For instance, from the Golgi to the endoplasmic reticulum (ER) and from the Golgi to the PM along the biosynthetic-secretory pathway, or from the PM to the vacuole through the endosome through the endocytic pathway. Interestingly, vacuole intrinsic vesicular transport (Echeverría, 2000) and transport vesicles in other organelles like chloroplasts (Karim and Aronsson, 2014) and mitochondria (Yamashita et al., 2016) have also been observed. Besides structural and functional similarities with mammalian and yeast vesicles and vesicle mediated trafficking, plant cells show considerable differences too. One of the peculiarities, for instance, is the mobile secretory vesicles cluster that carry cargo molecules to the extracellular space through the PM and cell plate (Toyooka et al., 2009). Moreover, the existence of COPII transport vesicles in highly vacuolated plant cells is argued along with the mechanism that drive membrane trafficking between the ER and the Golgi apparatus (Robinson et al., 2015). In plants, the existence, secretion and role of extracellular vesicle (EVs) are much less understood that those of mammalian cells and biological fluids which are well characterized (An et al., 2007; Regente et al., 2012; Rutter and Innes, 2017). In fact, there are only few studies reporting characterization of EVs isolated from the extracellular apoplastic fluids of Arabidopsis leaf (Rutter and Innes, 2017) and sunflower seeds (Regente et al., 2009).

Recently, nanometer-sized membrane vesicles have been isolated from complex matrixes, such as homogenized roots of ginger and carrot (Mu et al., 2014) and the fruit juices of grape (Ju et al., 2013), grapefruit (Wang et al., 2013), and lemon (Raimondo et al., 2015) using the gradient ultracentrifugation (UC) method commonly used for the isolation of mammalian EVs. It has been shown by physiochemical methods (like electron microscopy, EM and dynamic light scattering, DLS) and by molecular characterization (proteins, lipids and RNAs cargo) that plant-derived nanoparticles are indeed vesicles. Bulk vesicle preparations form plants show similarities with that of mammalian EVs, including a broad size-distribution (between 20 and 500 nm in diameter) and buoyant density. Bulk plant vesicles have been separated into a number of different vesicle populations by density gradient UC. Isolated nanovesicles have been studied for their cellular uptake, cytotoxicity, biological and therapeutic effects both *in vitro* (Ju et al., 2013; Raimondo et al., 2015) and *in vivo* (Ju et al., 2013; Raimondo et al., 2015; Wang et al., 2013). Mu *et al*. recently suggested that plant derived nanoparticles may have an active role not only in intra- and intercellular cargo delivery but also, when ingested, in interspecies communication between plant and mammalian cells which opens up new avenues for further exploitation. Vesicles from grape were shown to be efficiently taken up by intestinal stem cells and to induce tissue remodeling and protection against colitis (Ju et al., 2013). Besides their direct use, plant nanovesicles are being exploited as a novel technology platform for loading and delivery of exogenous nutraceuticals and drugs too (Wang et al., 2013). For example, grapefruit-derived nanovectors have been successfully used to load and deliver a variety of therapeutic agents including chemotherapeutic drugs, DNA expression vectors, siRNAs and antibodies to intestinal macrophages (Wang, B. et al., 2014). Edible plant derived nanovesicles in comparison with current drug delivery nanosystems, such as liposomes, gold nanoparticles, carbon nanotubes, polymeric nanoparticles, dendrimers, etc. show advantages in terms of naturally non-toxic origin, cellular uptake, biocompatibility and reduced clearance rates (Zhang et al., 2016). Moreover, large-scale production of plant nanovesicles is much more feasible than isolation of mammalian cell-derived EVs from culture media. Hence translation of plant-derived vesicles hold great promise for potential pharmaceutical, nutraceutical and cosmeceutical applications. In spite of their importance, at present we have only limited knowledge about the molecular composition and heterogeneity of the various vesicle populations isolated from plant tissues.

Citrus is one of the most important fruit crops in the world. Consumption of orange juice for example is associated with improved diet quality and positive health outcomes, including lower total cholesterol and low density lipoprotein levels. *Citrus limon* juice-derived nanovesicles were shown to inhibit cancer cell proliferation *in vitro* and *in vivo* through the activation of TRAIL-mediated apoptosis (Raimondo et al., 2015; Raimondo et al., 2018). Citrus species will likely become a major source of nanovectors in food and healthcare biotechnology. In this study, we isolated vesicle-like organelles from the fruit juice of four *Citrus* species: *C. sinensis, C. limon, C. paradisi* and *C. aurantium*. Micro-(MV) and nanovesicle (NV) fractions were separated, purified and their protein cargo were characterized and compared using a label free quantitative shotgun proteomics. The main aim was to identify elements of the highly complex vesicular transport network and to shed light on the physiological and biological roles of the proteins especially enzymes present. A bioinformatics approach, including a comparative analysis with a recently published ‘core set’ of predicted vesicular transport proteins of *A. thaliana* and yeast corresponding to 240 factors (Paul et al., 2014) and proteins present in EVs databases (Kim et al., 2015) were used as baits to identify putative vesicular transport and EVs related components in citrus vesicles.

## 2. Material and methods

### 2.1 Plant material and vesicle isolation

Fruits of four different Citrus species, sweet orange (*C. sinensis*), lemon (*C. limon*), grapefruit (*C. paradisi*) and bitter orange (*C. aurantium*) grown without any pre- and postharvest treatments were collected at maturity stage 3 from local gardens in Naples, Italy in July 2017. 5-10 pieces of fresh fruits were squeezed to obtain approximately 500 mL of fruit juice. Protease inhibitor cocktail containing cOmplete ultra tablets EDTA free (Roche, Mannheim, Germany) prepared according to manufacturer instruction, Leupeptine (0.25 mL, 1 mg/mL), 100 mM Phenylmethylsulfonyl fluoride (PMSF) (1.25 mL) and 1 M sodium azide (0.8 mL) were added to each samples. Vesicles were isolated in four parallel experiments by differential ultracentrifugation following the procedure described by Stanly *et al*. (Stanly et al., 2016). Briefly, low-velocity centrifugation steps were performed at 400 × *g* and 1000 × *g* for 20 min at 22 °C to eliminate cell debris, and at 15,000 × *g* for 20 min at 22 °C to collect the fraction enriched in microvesicles. The supernatant was ultracentrifuged at 150,000 × *g* for 60 min at 4 °C using a 70Ti Beckman rotor and the resulting pellet was set aside for nanovesicle purification. To remove co-purifying pectins, pellets obtained after the 15,000 × *g* and 150,000 × *g* centrifugation steps were solubilized in 50 mM Tris-HCl pH 8.6 and re-centrifuged two more times using the same centrifugal conditions. Finally, the Tris-HCl buffer was exchanged to PBS. Protein concentrations were measured by micro BCA assay (Thermo Scientific, Rockford, IL USA) using Nanodrop 2000 (Thermo Fisher Scientific Inc., Waltham, MA, USA). Samples were used fresh for subsequent characterization and proteomics experiments and then conserved at -80 °C.

### 2.2 Physiochemical characterization of different vesicle populations

#### 2.2.1 Transmission electron microscopy (TEM)

Samples were diluted to 1 μg/μL concentration using 0.1 M PBS pH 7.6 and five microliter aliquots were used for TEM analysis. The samples were deposited onto formvar and carbon coated 300 mesh copper grids for one minute. The preparation droplets were removed and the grids were dried. The samples were negatively stained with 2 % (w/v) aqueous uranyl acetate. TEM images were acquired by a Jeol JEM 1011 electron microscope operating at 60 kV and mounted with a Morada CCD camera (Olympus Soft Imaging Solutions).

#### 2.2.2 Dynamic Light Scattering (DLS)

Vesicle size distribution was measured by dynamic light scattering (DLS) using a Zetasizer Ver. 7.01, Malvern Instrument (Malvern, UK). Each sample was dispersed in deionized water and the intensity of the scattered light was measured with a detector at 90° angle at room temperature. Mean diameter and size distribution were the mean of three measures.

#### 2.2.3 Proteomics and data analysis

##### Protein profiling by SDS-PA GE

The quality of vesicle samples was controlled using sodium dodecyl sulfate polyacrylamide gel electrophoresis (SDS-PAGE). Samples (20 μg protein) were electrophoretically separated under reducing conditions on a precast Novex Bolt 4-12 % Bis-Tris Plus gel using Bolt MOPS SDS running buffer (Life Technologies, Carlsbad, CA, USA) according to the manufacturer’s instructions and stained with colloidal Coomassie blue.

##### Shotgun proteomics analysis

For *in-solution* proteomics analysis, samples (100 μg protein measured by micro BCA assay) were solubilized in 0.2 % RapiGest SF (Waters Corp., Milford, MA, USA) and vesicles were lysed using five freeze-thaw cycles under sonication in dry ice with ethanol. Proteins were reduced using 5 mM DL-Dithiothreitol (Sigma-Aldrich, Saint Louis, MO, USA,), alkylated using 15 mM Iodoacetamide (Sigma-Aldrich, Saint Louis, MO, USA), and digested using MS grade trypsin (Pierce, Thermo Sci. Rockford, IL, USA) as described previously (Stanly et al., 2016). Samples were vacuum dried and solubilized in 5 % acetonitrile and 0.5 % formic acid (LC-MS grade, Thermo Sci. Rockford, IL, USA) prior to nano-HPLC-MS/MS analysis.

##### nanoLC-ESI-MS/MS

Shotgun proteomics analysis was carried out on 1 μg of tryptic digest using a Dionex Ultimate 3000 nanoRSLC (Dionex, Sunnyvale, Ca, USA) coupled to a Bruker Maxis II mass spectrometer (Bruker Daltonics GmbH, Bremen, Germany) via CaptiveSpray nanobooster ionsource. Peptides were desalted on an Acclaim PepMap100 C-18 trap column (100 μm × 20 mm, Thermo Scientific, Sunnyvale, CA, USA) using 0.1 % TFA for 8 minutes at a flow rate of 5 μL/min and separated on the ACQUITY UPLC M-Class Peptide BEH C18 column (130 Å, 1.7 μm, 75 μm × 250 mm, Waters, Milford, MA, USA) at 300 nl/min flow rate, 48 °C column temperature. Solvent A was 0.1 % formic acid, solvent B was acetonitrile, 0.1 % formic acid and a linear gradient from 4 % B to 50 % B in 90 minutes was used. Mass spectrometer was operated in the data dependent mode using a fix cycle time of 2.5 sec. MS spectra was acquired at 3 Hz, while MS/MS spectra at 4 or 16 Hz depending on the intensity of the precursor ion. Singly charged species were excluded from the anaylsis.

##### Protein identification, quantitation and statistical analysis

Raw data files were processed using the Compass DataAnalysis software (Bruker, Bremen, Germany). Proteins were searched using the Mascot software v.2.5 (Matrix Science, London,UK).against the NCBI *Citrus sinensis* database (88176 entries). The search criteria were the following: 7 ppm precursor and 0.05 Da fragment mass tolerance. Two miscleavages were allowed, and the following modifications were set: Carbamidomethylation on cysteine as fixed modification, while methionine oxidation and asparagine and glutamin deamidation as variable modifications. Protein identifications were validated by the Percolator algorithm (Käll et al., 2007), false discovery rate was < 1%. The default peak-picking settings were used to process the raw MS files in MaxQuant (Cox and Mann, 2008) version 1.5.3.30 for label free quantitation. Peptide identifications were performed within MaxQuant using its in-built Andromeda search engine. Heat map was generated by XLStat statistical software ad-in in Microsoft Excel (Langella et al., 2013).

##### Bioinformatics

Functional annotation and data mining was performed using Blast2Go v.4.1.9 software including InterPro, enzyme codes, KEGG pathways, GO direct acyclic graphs (DAGs), and GOSlim functions (Conesa and Gotz, 2008). Identified proteins in FASTA format was used as input data. The Blastp-fast search algorithm was used via NCBI web service using taxonomy filter green plants (taxa: 33090, Viridiplantae), number of blast hits 20 and expectation value 1.0E-3. The InterPro domain searches were performed using the input FASTA files. The Blast hits of each protein sequence were mapped with Gene Ontology (GO) terms deposited in the GO database. Enzyme Code (EC) EC and KEGG annotations were performed on hits resulted in GO annotation. Venn diagrams were prepared using FunRich open access standalone software (Pathan et al., 2015). Orthology assignment and clusters for orthologous groups (COG) annotation were performed by EggNOg Mapper version 4.5.1 (Huerta-Cepas et al., 2016). Functional annotation in EggNOG is based on fast orthology assignments using precomputed eggNOG clusters and phylogenies. EggNOG OGs were used to compare protein datasets between different taxa and EVPedia (Kim et al., 2015) deposited OGs. For the identification of vesicular transport related proteins in citrus vesicles, Sequence Bulk Download and Analysis tool (https://www.arabidopsis.org/tools/bulk/sequences/index.jsp) was used to retrieve the Arabidopsis Genome Initiative (AGI) locus identifiers of the recently published Arabidopsis core-set (Paul et al., 2014) into FASTA format using all gene model/splice form settings. FASTA file then was imported and analyzed by Blast2Go and EggNOG software.

## 3. Results and Discussion

### 3.1 Characterization of micro- and nanovesicles fractions of citrus-derived vesicles

The citrus juice sac cells originate from anticlinal and periclinal divisions of epidermal and subepidermal cells of the endocarp. The mature juice sac consists of an external layer of elongated epidermal and hypodermal cells which enclose large highly vacuolated and thin-walled juice cells (Burns et al., 1992). Juice of citrus fruit contains a mixture of disrupted cells and cellular fluids. After removal of cells, cellular debris and large organelles, vesicle populations of the juices were separated into MVs and NVs enriched fractions using differential UC. Here, according to current EVs nomenclature vesicles were distinguished depending on their rough diameter size: MVs from 100 to 1,000 nanometers (nm) and NVs from 50 to 100 nm. The quantity of vesicles was estimated based on protein concentration. Different preparations contained variable amounts of proteins. MVs: 1445 mg kg^-1^ of grapefruit, 985 mg kg^-1^ of orange, 187 mg kg^-1^ of bitter orange and 94 mg kg^-1^ of lemon. NVs: 105 mg kg^-1^ of lemon, 52 mg kg^-1^ of orange, 32 mg kg^-1^ of grapefruit and 25 mg kg^-1^ of bitter orange.

The MVs and NVs fractions were characterized by particle size distribution and morphology using DLS and TEM analysis (Figure 1 A and B) and by profiling of their respective protein biocargo using SDS-PAGE (Figure 1 C) and shotgun proteomics. Particles ranged in size from approximately 265 (grapefruit) to 950 nm (orange and bitter orange) in diameter were detected in the MV rich fraction by DLS. Vesicles found in the NV rich fractions showed a 2-mode distribution: the size of smaller particle populations ranged approximatively from 50 to 80 nm, while bigger vesicle groups measured from 350 to 700 nm (Figure 1 and Supplementary Figure 1-3). TEM confirmed that all the samples contained intact vesicles. Nucleus or other cellular compartments were not detected in these fractions. In accordance with the DLS findings, TEM analysis shows the presence of large particles in the MV fractions and similarly to DLS analysis, distinguish two particle populations in the NV enriched fractions as shown in Figure 1 B and Supplementary Figure 1-3. SDS-PAGE-based protein profiles of both MV and NV fractions of different citrus species are highly complex (Figure 1 C), an aspect that resembles to that of EVs isolated from mammalian cells. Protein patterns of the four different citrus species were similar to each other, but the SDS profiles of MVs and NVs were markedly different.

### 3.2 The protein cargo of different citrus vesicles

The protein cargo of vesicles is directly implicated in various events, like vesicle formation, curvature shaping, binding, as well as transfer of biological function to the membrane cell or donor organelle. The protein cargo of vesicles is packaged inside the lumen or it is part of the vesicle membrane (integral membrane proteins). In this study we applied organelle proteomics to identify the protein cargos of fruit sac cells-derived vesicle populations from four citrus species with the aim to compare them with existing datasets. Proteins were identified against a subset of the NCBI protein sequence database limited to *C. sinensis* taxonomy group and their expression levels were compared by means of label-free quantitative proteomics. *C. sinensis* taxonomy was chosen due to the successful completion of the sweet orange genome sequencing project in 2012 (Wang, J. et al., 2014) and because it has the highest number of protein sequences amongst citrus species deposited in NCBI database. In each samples approximately 600-800 proteins were identified using the Mascot search engine (Supplementary Table 1 A and B). Label-free protein quantitation was performed by MaxQuant to determine relative protein expression levels (Supplementary Table 2A). In the 8 samples analysed we have quantified altogether 1700 proteins. The Venn diagrams (Figure 1 D) display the numbers of unique and common proteins quantified in each sample. Altogether, there were 577 proteins in the MV and 440 proteins in the NV fractions commonly expressed by all the four species, and 389 proteins were mutual among the 8 samples analyzed. Orthologous groups (OGs) of each identified protein were predicted using EggNOG mapper (Huerta-Cepas et al., 2016). OGs were used to compare our data to other taxonomically different datasets, and in particular to that of mammalian EVs related proteins deposited in EVPedia database (Kim et al., 2015). Significant OG hits searched against the OG accession codes (23618 hits) published in EVPedia resulted in 769 overlapping OGs (Figure 2A) and 1408 associated protein hits. This indicates that a high percentage (87%) of the protein cargo of citrus vesicles mapped by EggNOG (i.e. 1408 out of the 1694 proteins listed in Supplementary Table 2B) were aligned to EVPedia. This set of proteins was considered to be relevant to the extracellular vesicle component (i.e. apoplastic vesicles) of NV and MV fractions. Similarly, we compared the predicted OGs associated to the protein cargo of the NV fractions of *C. limon* and *C. paradisi* to datasets from previous studies (Raimondo et al., 2015; Wang, B. et al., 2014) (Figure 2B). It should be noted that both studies have used gradient ultracentrifugation for the purification of the NVs after differential centrifugation yielding more homogeneous vesicle populations comparing to the NV fractions studied here. 57% of the OGs were found to be common in *C. limon* associated to 65% of the proteins identified by Raimondo *et al* and 35% of the OGs associated to 42% of the proteins identified by Wang *et al*, in the considerably smaller data set encompassing 98 proteins matched by EggNOG, in *C. paradisi* (Figure 2B and Supplementary Table 2B). These data indicate that beside the common set of proteins, the vesicular protein cargo of citrus is characterized by distinct features depending on, amongst other factors, the species, variants, samples, type of vesicle population as well as sample preparation method.

### 3.3 Candidate protein markers for citrus vesicles

EV research is facilitated by highly expressed EV marker proteins, like the tetraspanin family (CD9, CD63 and CD81), cytosolic proteins (such as heat shock 70 kDa protein, HSP70) and tumor susceptibility gene 101 protein (TSG101) which are frequently used in immunoblotting experiments and are useful to prove the vesicle character of a sample. Similar markers have not yet been described for plant-derived vesicles. In this study, we analyzed the 20 top ranking proteins commonly identified in MVs and NVs fractions of juice sac cells to find marker candidates for these types of vesicles. Accession number of cluster of orthologous groups (COG) were determined and compared to that of “Top100 + EV marker” published in EVPedia database (http://student4.postech.ac.kr/evpedia2_xe/xe/) using identification count. Identification count is a number that reflects the frequency of identification of a given protein and its orthologues in EV datasets. We found that orthologues of most of the top ranking proteins in our data sets (18 out of 20 in both MV and NV fractions) have previously been identified in EVs (Table 1). This is in accordance with previous findings and shows that vesicles from different sources have a recurrent set of membrane and cytosolic proteins. In our set these are Enolase, HSP70, HSP80, 14-3-3 family proteins, glyceraldehyde-3-phosphate dehydrogenase (G3PD) and fructose-bisphosphate aldolase 6 (FBA6). These top-ranking proteins have very high EVPedia identification codes (higher than 400) and are amongst the 23 most frequently occurring orthologues.

Two fascinating plant proteins, *Patellin-3-like* and *clathrin heavy chain*, were found to be highly expressed in all the citrus samples (Table 1 and Supplementary Table 3A). *Patellin-3-like* is the most abundant protein in the citrus MV (Table 1A) and the second most abundant in the NV fractions (Table 1B). Patellins (PTLs) have been characterized in *Arabidopsis, zucchini* (*Cucurbita pepo*) and *Glycine max* (Sha et al., 2016) but not yet in citrus. PTL proteins share the typical domain structure that is a SEC14 lipid binding domain, a C-terminal Golgi dynamics domain (GOLD) and an acidic N-terminal domain. PTL member PTL3 is a plasma-membrane protein involved in vesicle/membrane-trafficking events associated with high cell division activity like cell plate formation during cytokinesis. PTL3 binds hydrophobic molecules such as phosphoinositides and promotes their transfer between the different cellular compartments. Most recently microarray-based approach demonstrated that PTLs play a fundamental role in cell polarity and patterning by regulating the auxin-induced PIN (Auxin efflux carrier family protein) relocation in *Arabidopsis thaliana* (Tejos et al., 2018). The presence of PTL in vesicular cargo is novel and its biological significance is worth further study. *Clathrins*, on the other hand, are well-known scaffold proteins that play a major role in the formation of clathrin coated vesicles (CCVs). CCV formation in mammals requires the assembly of a polyhedral cage composed of clathrin heteropolymers of heavy and light chains. CCVs mediate the vesicular transport from the trans-Golgi network/Early Endosome to the PM. Clathrin-mediated endocytosis (CME) as a major route of endocytosis in plant cells. The high abundance of *clathrin heavy chain-1* coat components in all of the samples and the presence of *clathrin light chain 1-like* protein in *C. paradisi* and *C. limon* samples indicates that the isolated citrus sac cell vesicles are predominantly clathrin coated.

Interestingly, aquaporin, a protein frequently detected in mammalian extracellular vesicles was detected only in the NV fraction (Supp. Table 1A and 2A). Recently aquaporins have been reported to play an important role in vesicle stability in case of plasma membrane vesicles of broccoli plant. (Martínez-Ballesta et al., 2018) Based on these observations, putative protein markers for plant vesicles are HSP70, HSP80, 14-3-3, G3PD and FBA6, PTL3 and clathrin proteins. On the other hand for plant nanovesicles a protein marker candidate is aquaporin. These proteins can facilitate the targeted functional and expression studies in the field of plant-derived vesicles.

### 3.4 Quantitative analysis of protein biocargo of citrus vesicles

A heat map chart in Figure 3 was generated using the MaxQuant output of the label-free shotgun analysis re-arranged according to clusters produced by hierarchical clustering. It shows the quantified proteins clustered in rows and the 8 citrus samples (MV and NV of 4 citrus species) in columns. By analyzing sample and protein dendrograms individually, it can be seen that the 8 samples are divided into two main groups (upper dendrogram). The clusters on the upper left and the upper right sides corresponds to the 4 MVs and the 4 NVs fractions, respectively. This shows that the vesicle cargo in different size vesicles result in a much larger difference, than that among the various citrus species. In the NV fractions *C. sinensis* and *C. paradisi* on the one hand, and *C. limon* with *C. aurantium* on the other hand show a large similarity. This corresponds well with philological relationship: *C. paradisi* is a hybrid of *C. sinensis* and *C. maxima*, while *C. limon* is a hybrid between *C. aurantium* and *C. medica*. In the left dendrogram proteins are divided into three major groups. The green and red large rectangles on the lower part of the map correspond to the first hierarchical cluster of proteins show a relatively high expression of this cluster of proteins in the MVs (green) and low expression in the NVs (red).

### 3.5 Heterogeneity of trafficking vesicles in citrus juice sac cells

Cellular vesicles are involved in the intra and inter cellular transport and communication events regulated by the secretory, recycling, endo- and exocytic pathways. Specialized proteins, for example, the components of endosomal sorting complexes required for transport (ESCRT), the Rab GTPases which are involved in regulation of vesicle formation and uncoating, and the SNAREs and tethering factors which aid the membrane fusion processes regulate these pathways. These proteins are located within the vesicle’s coat or recruited from the cytosol while some others are expressed on the membrane surface of the vesicle-interacting cell or organelle. Plants have been shown to possess a high number of putative orthologues and paralogous for known genes that encode for the respective coat proteins (Faini et al.), Rab GTPases (Flores et al., 2018), SNAREs (Lipka et al., 2007) and ESCORTs (Gao et al., 2014). Most vesicular transport related proteins in plants are predicted by bioinformatics and thus far only few of them have been experimentally identified. One such example is FREE1, which is a unique plant ESCRT component regulating multivesicular body protein sorting and plant growth in Arabidopsis (Gao et al., 2014). Given the importance of vesicular trafficking for physiological (Yun and Kwon, 2017) and pathological but also for environmentally triggered processes (like salinity, (Baral et al., 2015)), it is of prime importance to experimentally determine and characterize the expression of vesicular transport related proteins in plants. Here, we analysed the experimentally determined citrus juice sac cell related proteome using bioinformatics with the aim to identify vesicular trafficking machinery related proteins including vesicle coat proteins indicative of the type of trafficking vesicle (CCV, COPI and COPII). Proteins were indicated to be related to vesicular trafficking based on two different methods: 1) finding orthologues based on previously described vesicular transport proteins and 2) GO term enrichment analysis.

Recently, vesicular transport proteins of *A. thaliana* and yeast corresponding to ‘core-set’ of 240 factors (331 genes) has been used in an *in silico* study to detect putative proteins and group of (co-)orthologues in 14 plant species.(Paul et al., 2014) The ‘core-set’ contains 8 factors for the COPII, 16 for COPI, 18 for CCVs, 20 for Retromers and ESCRTs, 68 for Rab GTPases, 45 for Tethering factors and 65 for SNAREs. To determine putative vesicular transport related proteins and orthologue groups in the *C. sinensis* proteome, the Arabidopsis ‘core-set’ was translated into proteins using The Arabidopsis Information Resource (TAIR) database and used as a bait in B2Go blast search. 474 out of 489 Arabidopsis proteins yielded 1 to 20 blast hits and resulted altogether 1045 putative vesicular transport related proteins in *C. sinensis* (Supplementary Table 3A). Exploration of this set of proteins against the vesicular proteome of 4 citrus species identified in this work (Supplementary Table 1) resulted in 91 proteins (Supplementary Table 3B).

Additional putative vesicle related transport proteins of citrus were identified by extracting the proteins that are associated to GO terms “Transport) (GO:0006810) (Figure 4 A). Out of 299 proteins in our experimental data set associated with transport 96 were related to “vesicle-mediated transport” (GO:0016192). Comparing this with the set of proteins identified based on the ‘core-set’ of Arabidopsis (Figure 4 B) resulted in 119 putative vesicular related transport proteins (Supplementary Table 3B). Analysis of the dataset reveals the presence of proteins related to all the three main types of intracellular vesicles: CCV (8 proteins) and COPI (13 proteins) and COPII machineries related proteins (8 proteins). Coated vesicles (COPI, COPII and CCVs) are formed in a similar manner. The formation of vesicles is generally induced by a conformational change triggered by the exchange of guanosine diphosphate (GDP) to guanosine triphosphate (GDT) within a small GTPase. The GTPase recruits adaptor proteins, which in turn recruit coat proteins from the cytosol that polymerize to form the outer coat (Dodonova et al., 2015). The presence of specific coat proteins thus is indicative of the type of vesicle.

***COPI*** transport vesicles mediate retrograde transport from the Golgi to the ER and within the Golgi in the early secretory pathway. The key component of the COPI coat is a coatomer protein complex composed of seven subunits (α-, β-, β’-, γ-, δ-, ε- and ζ-COP) in eukaryotes. Several isoforms of α-, β-, β -, ε- and ζ- subunits, but not isoforms for γ-COP and δ-COP subunits have been identified in Arabidopsis genome. Here we identified seven isoforms of α subunit of the outer layer of the of the COPI coat (Supplementary Table 3B). Other subunits of the COPI coatomer complex were not identified. In addition, six ADP-ribosylation factors (Arfs) could also be identified. Arfs are members of the ARF family of GTP-binding proteins of the Ras superfamily implicated in the regulation of COPI coat assembly and disassembly. Beside triggering the polymerization of the coat proteins and budding of the vesicles growing evidence supports the structural presence of Arf1 within the coat in COPI (Dodonova et al., 2017).

***COPII*** intact spherical membrane vesicles are about 60–70 nm in size that carries protein cargo to the ER-Golgi intermediate compartment. The COPII polymerized coat includes five cytosolic proteins: Sar1, Sec23, Sec24, Sec13 and Sec31. The inner COPII coat is formed by the Sec23 and Sec24 heterodimers and the outer coat is constructed by Sec13 and Sec31 hetero-tetramers. In our experimental dataset homologues of all the five COPII coat proteins are present. Sar1 protein, a monomeric small GTPase regulates the assembly and disassembly of COPII coat and Sec12, of which homologue could also be identified in this work, is the guanine nucleotide exchange factor (GEF).

***CCVs*** are produced at the TGN, endosomes and plasma membrane and are involved in various post-Golgi trafficking routes and endocytosis. Clathrin-mediated endocytosis is a well-established process not only in animal but also in plant cells. The outer shell of the CCV is a cage of interlocking clathrin triskelions, a large heterohexameric protein complex composed of three heavy and three light chains. Clathrin coat proteins are not able to bind directly to the membranes therefore their connection is mediated by so called clathrin adaptors. The AP complexes mediate both the recruitment of clathrin to membranes and play a central role in membrane trafficking by packaging cargo proteins into nascent vesicles. Arabidopsis genome encodes all subunits (adaptins) of five APs (AP-1 to AP-5). In our dataset 8 CCVs factors have been identified including the clathrin light and heavy chains coat proteins and subunits of adaptins AP-1, AP-2, AP-3 and AP-4. AP-1 is located at the TGN and involved in vacuolar trafficking and secretion in non-dividing cells, and trafficking to the cell plate from the TGN in dividing cells. AP-2 is involved in endocytosis from the PM and the cell plate. The AP-3 beta adaptin that mediates the biogenesis and function of lytic vacuoles in Arabidopsis was also identified in the dataset. In this work we do not identified homologues subunits of AP-5.

Other vesicular transport related proteins revealed in this study are Rab GTPAse (25 proteins), SNARE (18 proteins), tethering factors (12 proteins), retromer and ESCRTs proteins (15 proteins). Rab GTPase serves as master regulators of membrane trafficking starting from the formation of the transport vesicle till to its fusion at the target membrane. Specific Rab GTPAse is associated with each organelles. Her we identified 10 proteins from the largest RAB group called RABA1 that regulates transport between TGN and plasma mebrane. They all are isoforms of RABA1f proteins which gene in Arabidopsis is mainly expressed in flowers’ pollen tube. SNAREs are membrane proteins that mediate fusion between vesicles and organelles to transport cargo molecules within the cells. Some SNAREs are localized in vesicles (called v-SNARE) while other SNAREs (called t-SNARE) are on the target organelle. Physically SNARE-SNARE interactions are thought to occur very selectively in the cells interaction between SNAREs localized in the target. Because selected members of the SNARE family proteins are localized in selected types of organelles.

Quantitative analysis of the 119 proteins potentially related to vesicle trafficking shows that the most abundant vesicle mediated transport related proteins when taking the average MaxQuant LFQ results of the 8 Citrus samples were RAB GTPases (37.3% of all transport related proteins), followed by CCVs (19.4%) and SNARE vesicles (14.5%). COPI and COPII related proteins were present in 7.5% and 4.0%, respectively. The top 30 most abundant transport related proteins were detected in at least 7 out of the 8 samples. Analyzing the transport related proteins of the MV and NV fraction separately some differences appear. SNARE proteins are less abundant in the MV fraction (9.2%) than in the NV fraction (16.4%).

### 3.6 Ubiquity of hydrolases and oxidoreductases in citrus vesicles

Due to their efficient cellular uptake and lack of adverse immune reaction in human plant-derived vesicles are growingly exploited as nanovectors for the delivery of exogenous bioactive materials (Wang, B. et al., 2014; Wang et al., 2013). Prior encapsulation, it is important to know the biological effects that unloaded vesicles trigger on their recipient cells. Natural plant-derived vesicles were shown to have significant biological effects, like inhibition of cell proliferation, apoptotic effect on cancer cell lines and induction of intestinal stem cells. (Ju et al., 2013; Raimondo et al., 2015; Zhuang et al., 2015) Amongst the potential effectors there are different enzymes that are commonly associated to the protein cargo of the vesicles. Enzymes present in vesicles being protected by the phospholipid membrane and has been shown to survive protein degradation events and reach recipient cells in their active forms. Upon their release they are prompt to catalyze a diverse array of biochemical reactions within the recipient cells. This prompted us to analyse the enzyme-content of citrus vesicles.

A high number of enzymes (482 proteins, Supplementary Table 4) were found by performing Enzyme class (EC) annotation of proteins identified in citrus vesicle samples. These belong to the 6 main enzyme classes (i.e. 1. oxidoreductases, 2. transferases, 3. hydrolases, 4. lyases, 5. isomerases, and 6. ligases). Figure 5 shows these enzyme classes are distributed in citrus vesicles. The two most representative enzyme classes accounting for more than the half of the enzymes identified (Supplementary Table 4) are class 3 hydrolyses and class 1 oxidoreductases. The role of fruit **hydrolases** in physiological processes like fruit ripening is well studied, thus their high expression in citrus juice sac cell derived vesicles was expected. Amongst the hydrolases, Adenosine triphosphatases (ATPases EC:3.6.1.3) acting on acid anhydrides are the most ubiquitous enzymes in the all the samples studied (Figure 6). We have identified 44 ATPases, including Vacuolar-type H + -ATPases (V-ATPases), calcium-, magnesium- and phospholipid-transporting ATPases (Supplementary Table 4). V-type ATPAse pumps protons across the plasma membrane and has function in acidification. ATP-driven proton pumps are cell membrane localized though they are also ubiquitous in intracellular organelles, such as endosomes, Golgi-derived vesicles, secretory vesicles, and vacuole. Recent studies proposed acidification-independent function of vesicular V-ATPases directly related to membrane fusion and exocytosis (Wang and Hiesinger, 2013) and thus their presence can give indications on the origin of EVs. Additionally, high number of hydrolases that act on peptide bonds (43 enzymes, EC 3.4 in Figure 5) were found, like the 20S core proteasome subunits. Proteasome subunits accumulate in some microvesicles and have been shown to that the proteasome is involved in the regulation of MV release (Tucher et al., 2018). We observed the accumulation of almost all the 20S complex units (proteasome alpha 1B, 2A, 3, 5, 6 and 7; proteasome beta 1, 2A, 3A, 4, 5, 6 and 7B). Other highly expressed hydrolases in citrus vesicles were pectinesterase, phospholipases, amylases, β galatosidases and adenosylhomocystein hydrolyse.

**Enzymatic antioxidants** are very important enzymes in protecting the cell from oxidative damage caused by reactive oxygen species (ROS) as by-products of redox reactions. Enzymatic antioxidants have important roles in scavenging ROS. Members of catalase (CAT), superoxide dismutase (SOD), peroxidase (POD), ascorbate peroxidase (APX) as well glutathione reductase (GPX) family enzymes were found to be highly expressed in all the studied vesicle preparations (Supp. Table 4). There is a growing body of evidence suggesting that oxidative stress plays a key role in the pathogenesis of numerous human diseases, such as cancer, autoimmune disorders, aging, rheumatoid arthritis, cardiovascular and neurodegenerative diseases. Given the importance of the management oxidative stress in disease there is a considerable interest in identifying novel enzymatic antioxidants. In this context, plant-made exogenous antioxidants of SODs, CATs or GPXs family enzymes readily packed into vesicles could have huge potential.

### 3.7 Conclusions

Vesicles preparations isolated from complex matrixes, such as fruit juice through ultracentrifugation and ultrafiltration steps contains highly heterogeneous populations of extracellular and intracellular membrane vesicles separated roughly by size and density. Study of citrus fruit sac cells-derived vesicles of four different citrus species show the presence of highly expressed HSP70, HSP80, 14-3-3, G3PD and FBA6, PTL3 and clathrin proteins in both MV and NV fractions. Aquaporin was found to be enriched mainly in the NV fraction. Based on specific characteristics of the composition of protein cargo, the heterogeneity of vesicles could be dissected into different vesicle subpopulations. A high percentage of isolated vesicles shows extracellular origin based on orthology search against proteins expressed in EVs of mammalian origin (Figure 2). In addition, different subpopulations of intracellular vesicles, such as COPI, COPII and CCVs related to vesicular transport could be distinguished and characterized in citrus vesicles (Figure 4). Enzymes were highly expressed in both MV and NV fractions (Figure 5), and most significantly different hydrolases (ATPases, pectinesterase, phospholipases, amylases, β galatosidases and adenosylhomocystein hydrolyse) and enzymatic antioxidants (SODs, CATs, PODs and GPXs) were identified (Figure 6). Characterization of protein cargo of plant-derived vesicles is important in their exploitation as potential vectors for cellular delivery.

## Acknowledgements

This work was supported by a research grant No. SAC.AD002.037 Accordo Bilaterale CNR/HAS NutriC@rgo project, 2016-2018 CUP B66D16000360005.

---

**Figure 1.** Characterization of microvesicles (MVs) and nanovesicles (NVs) containing fractions isolated from fruit juice of *C. paradise* (selected as representative species for this figure) using differential ultracentrifugation,. A.) Images show particle-size distributions measured using dynamic light scattering. B.) Representative images of the vesicles observed by transmission electron microscopy of *C. paradisi* show MVs (B1-2) after the low velocity centrifugation (15.000 × g) step and both MVs (arrows) and NVs (B3) after differential ultracentrifugation at 150.000 × *g*. High magnification of NV rich area is demonstrated in panel B4. The scale bar indicates 500 nm in panel B1-3 and 100 nm in panel B4. C.) SDS-PAGE gel image of proteins of MV and NV enriched fractions of the four citrus species (*C. sinensis, C.limon, C. paradisi*, and *C. aurantium*). D.) Venn diagram (generated by FunRich software (Pathan *et al.*, 2015a)) shows the number of identified proteins identified in each citrus species in the MV and NV enriched fractions.

**Figure 2.** Comparisons of Orthologous Groups (OGs) of proteins identified in citrus vesicles: **A)** Venn diagram showing all the OGs of proteins identified by MaxQuant in vesicle fractions of the four citrus species and compared with that of EVPedia deposited data. **B)** Venn diagram of OGs of proteins identified in the nanovesicles fractions of *C. limon* and *C. paradisi* in this work and compared to that reported by Raimondo *et al* (Raimondo *et al.*, 2015) and Wang *et al* (Wang *et al.*, 2014a).

**Figure 3.** Cluster heatmap of two different citrus vesicles isolated from four citrus species. The columns of the heatmap represent the proteins and the rows represent the MV and NV samples of *C. sinensis, C.limon, C. paradisi*, and *C. aurantium*. The heat map shows expression values mapped to a color gradient from low (green) to high expression (red).

**Figure 4.** Putative vesicle-mediated transport related proteins identified in the micro and nanovesicle fractions of *C. sinensis, C.limon, C. paradisi*, and *C. aurantium*. A) Combined graph showing the number of protein sequences, node scores of GO:0016192 term related to vesicle mediated transport and the beneath lining GO terms. B.) Venn diagram showing the numbers of predicted vesicle-mediated transport related proteins extracted by the experimental data set using GO Term.

**Figure 5.** Distribution of the six main enzyme classes over all protein sequences identified and quantified in citrus fruit juice sac cell-derived vesicles studied.

**Figure 6.** Distribution of the two highly expressed enzyme subclasses, hydrolases and oxidoreductases in the citrus fruit juice sac cell-derived vesicles studied.

**Table 1.** List of 20 top ranking proteins commonly identified in all in the A) microvesicles and B) nanovesicles containing fractions of the four citrus species. **GI** indicates the NCBI access number of the proteins; **Length** is the number of the amino acids; **COG** is for cluster of orthologous groups accession number obtained from EggNOG mapper; **Enzyme name** was retrieved from the InterPro search of Blast2Go; EVPedia indicates the frequency score published in EVPedia database.

## Supplementary data

**Supp. Figure 1.** Particle size distribution and transmission electron microscopy (TEM) images obtained on the microvesicle (MV) rich fraction (panel A) and nanovesicle (NV) rich fraction (panel B) of *Citrus aurantium*. TEM panels show MVs (upper panel), both MVs (arrows) and NVs (arrowheads) (lower panel). High magnification of NVs is demonstrated in the insertion of the lower panel. TEM and DLS images relative to NV and MV rich fractions isolated from *C. paradisi* are shown in Figure 1. The scale bar indicates 500 nm in the upper TEM panel and indicates 100 in the lower TEM panels.

**Supp. Figure 2.** Particle size distribution and transmission electron microscopy (TEM) images obtained on the microvesicle (MV) rich fraction (panel A) and nanovesicle (NV) rich fraction (panel B) of *Citrus limon*. TEM panels show MVs (upper panel), both MVs (arrows) and NVs (arrowheads) (lower panel). High magnification of NVs is demonstrated in the insertion of the lower panel. TEM and DLS images relative to NV and MV rich fractions isolated from *C. paradisi* are shown in Figure 1. The scale bar indicates 500 nm in the upper TEM panel and indicates 100 in the lower TEM panels.

**Supp. Figure 3.** Particle size distribution and transmission electron microscopy (TEM) images obtained on the microvesicle (MV) rich fraction (panel A) and nanovesicle (NV) rich fraction (panel B) of *Citrus sinensis*. TEM panels show MVs (upper panel), both MVs (arrows) and NVs (arrowheads) (lower panel). High magnification of NVs is demonstrated in the insertion of the lower panel. TEM and DLS images relative to NV and MV rich fractions isolated from *C. paradisi* are shown in Figure 1. The scale bar indicates 500 nm in the upper TEM panel and indicates 100 in the lower TEM panels.

**Supp. Table 1.** Identified proteins in the MVs and NVs fractions of 4 citrus species.

**Supp. Table 2 Protein biocargo of citrus vesicles. A)** Label-free quantitative proteomics analysis of micro- and nanovesicle fractions obtained from the fruit juices of 4 citrus species (*C. sinensis*, *C. limon*, *C. paradisi* and *C. aurantium*), using MaxQuant software. **B)** OGs related to the vesicle protein cargo of citrus vesicles determined by MaxQuant, and comparision with i.) EVPedia, ii.) the data obtained on *C. limon* by Raimondo *et al* (Raimondo *et al.*, 2015) and iii.) the data obtained on *C. paradisi* by Wang *et al* (Wang *et al.*, 2014a). EggNOGs OGs were determined by EggNOG mapper version 4.5.1. (Huerta-Cepas et al., 2016).

**Supp. Table 3 A)** Putative vesicular transport related proteins in *Citrus sinensis*. **B)** Potential vesicular transport related cargo proteins identified in citrus vesicles.

**Supp. Table 4 A)** Enzyme Codes and expression analysis and **B)** Kegg Pathway analysis of enzymes identified in 4 citrus vesicle samples.

## References

An, Q., van Bel, A.J., Huckelhoven, R., 2007. Do plant cells secrete exosomes derived from multivesicular bodies? Plant Signal Behav 2(1), 4–7.

Baral, A., Shruthi, K.S., Mathew, M.K., 2015. Vesicular trafficking and salinity responses in plants. IUBMB Life 67(9), 677–686.

Burns, J.K., Achor, D.S., Echeverria, E., 1992. Ultrastructural Studies on the Ontogeny of Grapefruit Juice Vesicles (Citrus paradisi Macf. CV Star Ruby). International Journal of Plant Sciences 153(1), 14–25.

Conesa, A., Gotz, S., 2008. Blast2GO: A comprehensive suite for functional analysis in plant genomics. Int J Plant Genomics 619832(10), 619832.

Cox, J., Mann, M., 2008. MaxQuant enables high peptide identification rates, individualized p.p.b.-range mass accuracies and proteome-wide protein quantification. Nat Biotechnol 26(12), 1367–1372.

Dodonova, S.O., Aderhold, P., Kopp, J., Ganeva, I., Rohling, S., Hagen, W.J.H., Sinning, I., Wieland, F., Briggs, J.A.G., 2017. 9A structure of the COPI coat reveals that the Arf1 GTPase occupies two contrasting molecular environments. Elife 16(6), 26691.

Dodonova, S.O., Diestelkoetter-Bachert, P., von Appen, A., Hagen, W.J.H., Beck, R., Beck, M., Wieland, F., Briggs, J.A.G., 2015. A structure of the COPI coat and the role of coat proteins in membrane vesicle assembly. Science 349(6244), 195.

Echeverria, E., 2000. Vesicle-Mediated Solute Transport between the Vacuole and the Plasma Membrane. Plant Physiology 123(4), 1217–1226.

Faini, M., Beck, R., Wieland, F.T., Briggs, J.A.G., Vesicle coats: structure, function, and general principles of assembly. Trends in Cell Biology 23(6), 279–288.

Flores, A.C., Via, V.D., Savy, V., Villagra, U.M., Zanetti, M.E., Blanco, F., 2018. Comparative phylogenetic and expression analysis of small GTPases families in legume and non-legume plants. Plant Signaling & Behavior 13(2), e1432956.

Gao, C., Luo, M., Zhao, Q., Yang, R., Cui, Y., Zeng, Y., Xia, J., Jiang, L., 2014. A Unique Plant ESCRT Component, FREE1, Regulates Multivesicular Body Protein Sorting and Plant Growth. Current Biology 24(21), 2556–2563.

Huerta-Cepas, J., Forslund, K., Szklarczyk, D., Jensen, L.J., von Mering, C., Bork, P., 2016. Fast genome-wide functional annotation through orthology assignment by eggNOG-mapper. bioRxiv.

Hwang, I., Robinson, D.G., 2009. Transport vesicle formation in plant cells. Current Opinion in Plant Biology 12(6), 660–669.

Ju, S., Mu, J., Dokland, T., Zhuang, X., Wang, Q., Jiang, H., Xiang, X., Deng, Z.B., Wang, B., Zhang, L., Roth, M., Welti, R., Mobley, J., Jun, Y., Miller, D., Zhang, H.G., 2013. Grape exosome-like nanoparticles induce intestinal stem cells and protect mice from DSS-induced colitis. Mol Ther 21(7), 1345–1357.

Käll, L., Canterbury, J.D., Weston, J., Noble, W.S., MacCoss, M.J., 2007. Semi-supervised learning for peptide identification from shotgun proteomics datasets. Nature Methods 4, 923.

Karim, S., Aronsson, H., 2014. The puzzle of chloroplast vesicle transport – involvement of GTPases. Frontiers in Plant Science 5, 472.

Kim, D.K., Lee, J., Kim, S.R., Choi, D.S., Yoon, Y.J., Kim, J.H., Go, G., Nhung, D., Hong, K., Jang, S.C., Kim, S.H., Park, K.S., Kim, O.Y., Park, H.T., Seo, J.H., Aikawa, E., Baj-Krzyworzeka, M., van Balkom, B.W., Belting, M., Blanc, L., Bond, V., Bongiovanni, A., Borras, F.E., Buee, L., Buzas, E.I., Cheng, L., Clayton, A., Cocucci, E., Dela Cruz, C.S., Desiderio, D.M., Di Vizio, D., Ekstrom, K., Falcon-Perez, J.M., Gardiner, C., Giebel, B., Greening, D.W., Gross, J.C., Gupta, D., Hendrix, A., Hill, A.F., Hill, M.M., Nolte-’t Hoen, E., Hwang do, W., Inal, J., Jagannadham, M.V., Jayachandran, M., Jee, Y.K., Jorgensen, M., Kim, K.P., Kim, Y.K., Kislinger, T., Lasser, C., Lee, D.S., Lee, H., van Leeuwen, J., Lener, T., Liu, M.L., Lotvall, J., Marcilla, A., Mathivanan, S., Moller, A., Morhayim, J., Mullier, F., Nazarenko, I., Nieuwland, R., Nunes, D.N., Pang, K., Park, J., Patel, T., Pocsfalvi, G., Del Portillo, H., Putz, U., Ramirez, M.I., Rodrigues, M.L., Roh, T.Y., Royo, F., Sahoo, S., Schiffelers, R., Sharma, S., Siljander, P., Simpson, R.J., Soekmadji, C., Stahl, P., Stensballe, A., Stepien, E., Tahara, H., Trummer, A., Valadi, H., Vella, L.J., Wai, S.N., Witwer, K., Yanez-Mo, M., Youn, H., Zeidler, R., Gho, Y.S., 2015. EVpedia: a community web portal for extracellular vesicles research. Bioinformatics 31(6), 933–939.

Langella, O., Valot, B., Jacob, D., Balliau, T., Flores, R., Hoogland, C., Joets, J., Zivy, M., 2013. Management and dissemination of MS proteomic data with PROTICdb: example of a quantitative comparison between methods of protein extraction. Proteomics 13(9), 1457–1466.

Lipka, V., Kwon, C., Panstruga, R., 2007. SNARE-ware: the role of SNARE-domain proteins in plant biology. Annu Rev Cell Dev Biol 23, 147–174.

Martínez-Ballesta, M.d.C., García-Gomez, P., Yepes-Molina, L., Guarnizo, A.L., Teruel, J.A., Carvajal, M., 2018. Plasma membrane aquaporins mediates vesicle stability in broccoli. PLoS One 13(2), e0192422.

Mu, J., Zhuang, X., Wang, Q., Jiang, H., Deng, Z.B., Wang, B., Zhang, L., Kakar, S., Jun, Y., Miller, D., Zhang, H.G., 2014. Interspecies communication between plant and mouse gut host cells through edible plant derived exosome-like nanoparticles. Mol Nutr Food Res 58(7), 1561–1573.

Pathan, M., Keerthikumar, S., Ang, C.S., Gangoda, L., Quek, C.Y., Williamson, N.A., Mouradov, D., Sieber, O.M., Simpson, R.J., Salim, A., Bacic, A., Hill, A.F., Stroud, D.A., Ryan, M.T., Agbinya, J.I., Mariadason, J.M., Burgess, A.W., Mathivanan, S., 2015. FunRich: An open access standalone functional enrichment and interaction network analysis tool. Proteomics 15(15), 2597–2601.

Paul, P., Simm, S., Mirus, O., Scharf, K.D., Fragkostefanakis, S., Schleiff, E., 2014. The complexity of vesicle transport factors in plants examined by orthology search. PLoS One 9(5), e97745.

Raimondo, S., Naselli, F., Fontana, S., Monteleone, F., Lo Dico, A., Saieva, L., Zito, G., Flugy, A., Manno, M., Di Bella, M.A., De Leo, G., Alessandro, R., 2015. Citrus limon-derived nanovesicles inhibit cancer cell proliferation and suppress CML xenograft growth by inducing TRAIL-mediated cell death. Oncotarget 6(23), 19514–19527.

Raimondo, S., Saieva, L., Cristaldi, M., Monteleone, F., Fontana, S., Alessandro, R., 2018. Label-free quantitative proteomic profiling of colon cancer cells identifies acetyl-CoA carboxylase alpha as antitumor target of Citrus limon-derived nanovesicles. Journal of Proteomics 173, 1–11.

Regente, M., Corti-Monzón, G., Maldonado, A.M., Pinedo, M., Jorrín, J., de la Canal, L., 2009. Vesicular fractions of sunflower apoplastic fluids are associated with potential exosome marker proteins. FEBS Letters 583(20), 3363–3366.

Regente, M., Pinedo, M., Elizalde, M., de la Canal, L., 2012. Apoplastic exosome-like vesicles: a new way of protein secretion in plants? Plant Signal Behav 7(5), 544–546.

Robinson, D.G., Brandizzi, F., Hawes, C., Nakano, A., 2015. Vesicles versus Tubes: Is Endoplasmic Reticulum-Golgi Transport in Plants Fundamentally Different from Other Eukaryotes? Plant Physiology 168(2), 393–406.

Rutter, B.D., Innes, R.W., 2017. Extracellular Vesicles Isolated from the Leaf Apoplast Carry Stress-Response Proteins. Plant Physiology 173(1), 728–741.

Sha, A., Qi, Y., Shan, Z., Chen, H., Yang, Z., Qiu, D., Zhou, X., Chen, Y., Tang, J., 2016. Identifying patellin-like genes in Glycine max and elucidating their response to phosphorus starvation. Acta Physiologiae Plantarum 38(6), 138.

Stanly, C., Fiume, I., Capasso, G., Pocsfalvi, G., 2016. Isolation of Exosome-Like Vesicles from Plants by Ultracentrifugation on Sucrose/Deuterium Oxide (D2O) Density Cushions, in: Pompa, A., De Marchis, F. (Eds.), Unconventional Protein Secretion: Methods and Protocols. Springer New York, New York, NY, pp. 259–269.

Tejos, R., Rodriguez-Furlan, C., Adamowski, M., Sauer, M., Norambuena, L., Friml, J., 2018. PATELLINS are regulators of auxin-mediated PIN1 relocation and plant development in Arabidopsis thaliana. J Cell Sci 131(2), 204198.

Toyooka, K., Goto, Y., Asatsuma, S., Koizumi, M., Mitsui, T., Matsuoka, K., 2009. A Mobile Secretory Vesicle Cluster Involved in Mass Transport from the Golgi to the Plant Cell Exterior. The Plant Cell 21(4), 1212–1229.

Tucher, C., Bode, K., Schiller, P., Claβen, L., Birr, C., Souto-Carneiro, M.M., Blank, N., Lorenz, H.-M., Schiller, M., 2018. Extracellular Vesicle Subtypes Released From Activated or Apoptotic T-Lymphocytes Carry a Specific and Stimulus-Dependent Protein Cargo. Frontiers in Immunology 9, 534.

van de Meene, A.M.L., Doblin, M.S., Bacic, A., 2017. The plant secretory pathway seen through the lens of the cell wall. Protoplasma 254(1), 75–94.

Wang, B., Zhuang, X., Deng, Z.-B., Jiang, H., Mu, J., Wang, Q., Xiang, X., Guo, H., Zhang, L., Dryden, G., Yan, J., Miller, D., Zhang, H.-G., 2014. Targeted Drug Delivery to Intestinal Macrophages by Bioactive Nanovesicles Released from Grapefruit. Molecular Therapy 22(3), 522–534.

Wang, D., Hiesinger, P.R., 2013. The vesicular ATPase: A missing link between acidification and exocytosis. The Journal of Cell Biology 203(2), 171.

Wang, J., Chen, D., Lei, Y., Chang, J.-W., Hao, B.-H., Xing, F., Li, S., Xu, Q., Deng, X.-X., Chen, L.-L., 2014. Citrus sinensis Annotation Project (CAP): A Comprehensive Database for Sweet Orange Genome. PLoS One 9(1), e87723.

Wang, Q., Zhuang, X., Mu, J., Deng, Z.-B., Jiang, H., Zhang, L., Xiang, X., Wang, B., Yan, J., Miller, D., Zhang, H.-G., 2013. Delivery of therapeutic agents by nanoparticles made of grapefruit-derived lipids. Nat Commun 4.

Yamashita, A., Fujimoto, M., Katayama, K., Yamaoka, S., Tsutsumi, N., Arimura, S., 2016. Formation of Mitochondrial Outer Membrane Derived Protrusions and Vesicles in Arabidopsis thaliana. PLoS One 11(1).

Yun, H.S., Kwon, C., 2017. Vesicle trafficking in plant immunity. Current Opinion in Plant Biology 40, 34–42.

Zhang, M., Viennois, E., Xu, C., Merlin, D., 2016. Plant derived edible nanoparticles as a new therapeutic approach against diseases. Tissue Barriers 4(2), e1134415.

Zhuang, X., Deng, Z.B., Mu, J., Zhang, L., Yan, J., Miller, D., Feng, W., McClain, C.J., Zhang, H.G., 2015. Ginger-derived nanoparticles protect against alcohol-induced liver damage. Journal of extracellular vesicles 4, 28713.

